# Temporal distribution of cetaceans south of Pico Island (Azores, Portugal) - data obtained during whale-watching 2020

**DOI:** 10.1101/2022.06.07.495087

**Authors:** Peter Zahn, Lynn Kulike, Armin Bloechl

**Affiliations:** Institut of Biology and Chemistry, University of Hildesheim; Whale-watching guide; Institute for Terrestrial and Aquatic Wildlife Research (ITAW), University of Veterinary Medicine Hannover, Foundation

## Abstract

Land- and boat-based surveys were conducted to collect data during whale watching excursions in July/August and October 2020. Occurrence and distribution of cetaceans south of Pico Island (Azores) were determined. Ten species were sighted, 63 schools of Delphinids and 96 individuals of large whales. The most frequently sighted species were: Sperm whale (*Physeter macrocephalus*), Sei whale (*Balaenoptera borealis*), Atlantic spotted dolphin (*Stenella frontalis*), Common bottlenose dolphin (*Tursiops truncatus*), and Risso’s dolphin (*Grampus griseus*). An unusual high numbers of sightings of Sei whale in August and especially in October was noticed, and is not mentioned in the literature so far.

## Introduction

Places that were once popular for whaling are nowadays often used for whale-watching. The archipelago of the Azores developed to a hotspot of whale-watching since Espaço Talassa began as the first operator in the early 1990s in Lajes do Pico on the island of Pico. Tourism is growing rapidly and also whale-watching activities (Visser et al., 2011b). The Azores archipelago is a destination for nature tourism and one third of the guests practice whale-watching by boat. A wide variety of species can be observed here. 28 different species have been recorded in the Azores so far (Silva et al., 2014).

Information on residency, distribution, and abundance of cetaceans in this region remains limited (Cechetti et al., 2018). The knowledge comes mostly from a series of large-scale international surveys. This surveys are costly and surveying large areas of offshore waters comprise a lot of logistic and operational difficulties. The use of other data sets constitutes a valuable alternative for investigating how cetaceans use these areas (Silva et al., 2014). Whale-watching activities offer a source of valuable data and funding for cetacean research.

The data collected on marine life during whale-watching may contribute crucial data for understanding occurrence of cetaceans around the island of Pico and other Azorean islands (Bron et al., 2019). Precise knowledge of the distribution of the species is important for efficient protection of whales. The study aimed to gain some data of the distribution of the whales species observed in the waters of the south coast of Pico Island.

## Material and Method

In this study, data were collected in the Azores, an archipelago composed of nine volcanic islands in the North Atlantic Ocean. The survey (one land-based, 22 boat-based) took place 14 days in 2020, from July 24^th^ to August 2^nd^ and October 6^th^ to October 10^th^. Data were collected during commercial trips with the whale-watching company Espaço Talassa, with its homeport in Lajes do Pico, in the waters of the south coast of the island of Pico.

The excursion consisted of a group of 12 people in altogether, students and lecturers. The boat surveys were led by two skippers that represents the maximum capacity of 14 participants for one whale-watching boat. From this it follows that the number of trained observers were at least three. Each boat tour took three hours and occurred in the morning or in the afternoon respectively. A de-briefing after every boat-tour was carried out. All sightings were reviewed and questions answered. A comparison with the notes made on board was performed. Data collection was restricted. No tracked sail routes. Recorded number of boat tours are 22. Rough indicator of effort by determining the number of species encountered per trip.

## Results

During the study, in July/August and October 2020, a total of 10 whale species were observed (Tab. 1). 3.9 species were encountered per trip. In relation to their feeding ecology two main groups are to distinguish. Four teutophagous and deep diving species, *Physeter macrocephalus, Mesoplodon bidens, Globicephals macrorhynchus, and Grampus griseus*. The other six species feed on fish, squid or crustaceans, *Balaenoptera borealis, Delphinus delphis, Pseudorca crassidens, Stenella coeruleoalba, Stenella frontalis*, and *Tursiops truncatus* (Carwardine, 2020).

**Tab. 1:** Observed whale species with taxonomic classification.

| Order | Family | Scientific name | Common name |
| --- | --- | --- | --- |
| Mysticeti | Balaenopteridae | *Balaenoptera borealis* | Sei whale |
| Odontoceti | Physeteridae | *Physeter macrocephalus* | Sperm whale |
|  | Ziphiidae | *Mesoplodon bidens* | Sowerby's beaked whale |
|  | Delphinidae | *Delphinus delphis* | Short-beaked common dolphin |
|  |  | *Globicephala macrorhynchus* | Short-finned pilot whale |
|  |  | *Grampus griseus* | Risso's dolphin |
|  |  | *Pseudorca crassidens* | False killer whale |
|  |  | *Stenella coeruleoalba* | Striped dolphin |
|  |  | *Stenella frontalis* | Atlantic spotted dolphin |
|  |  | *Tursiops truncatus* | Common bottlenose dolphin |

Table 1 shows the sighted species of the four families. One species from each of the three families Balaenopteridae, Physeteridae and Ziphiidae were sighted. With six genera and seven species, members of the family Delphinidae were seen most frequently.

Only clearly identified species were counted. The individuals of the member of the first three families were countable due to their considerable body height and their relatively small group size. With a few exceptions this is not possible for the Delphinidae (Tab. 2). They are usually traveling in bigger schools.

**Tab. 2:** Number of sightings of whales.

| Species | No of sightings | No of Individuals | No of schools |
| --- | --- | --- | --- |
| *Balaenoptera borealis* | 20 | 35 |  |
| *Physeter macrocephalus* | 30 | 58 |  |
| *Mesoplodon bidens* | 1 | 3 |  |
| *Delphinus delphis* | 6 |  | 6 |
| *Globicephala macrorhynchus* | 8 |  | 8 |
| *Grampus griseus* | 12 |  | 12 |
| *Pseudorca crassidens* | 4 |  | 4 |
| *Stenella coeruleoalba* | 4 |  | 4 |
| *Stenella frontalis* | 17 | >1000 | 17 |
| *Tursiops truncatus* | 12 |  | 12 |

In this study, a total of 114 sightings (schools and individuals) of cetaceans were recorded (Tab. 2). A total of 96 individuals of Balaenopteridae, Physeteridae and Ziphiidae were seen. The family Delphinidae was represented by 63 schools. Repeated sightings are more than likely in both, individual whales and dolphin schools. Most frequently observed was *Physeter macrocephalus* with 30 sightings during the observation period. Followed by 20 recorded *Balaenoptera borealis*. Least observed species was *Mesoplodon bidens* with one sighting. Three individuals of the Beaked whales (Ziphiidae) were clearly identified as Sowerby’s beaked whale. Another 11 Ziphiids could not be identified definitely and were assigned to the genus Mesoplodon. Besides *Mesoplodon bidens* three more species of the genus Mesoplodon occur in the Azorean waters.

17 schools of *Stenella frontalis* were observed. *Grampus griseus* and *Tursiops truncatus* were among the frequently sighted cetacean species, with 12 sightings respectively. *Globicephala macrorhynchus* were represented with eight and *Delphinus delphis* with six schools respectively. *Stenella coeruleoalba* and *Pseudorca crassidens* were observed with four sightings each.

Sperm whale was observed every day (Fig. 1). The Sei whale was seen frequently, except on July 30^th^ and 31^st^. *Stenella frontalis* was observed often, except on July 25^th^, both days in August and on October 9^th^. *Delphinus delphis, Globicephala macrorhynchus* and Members of the genus Mesoplodon were recorded in July and August, *Mesoplodon bidens* was observed July 29^th^. *Stenella coeruleoalba* was only sighted in July. Common bottlenose dolphin was observed every day in August and October.

**Fig. 1:**
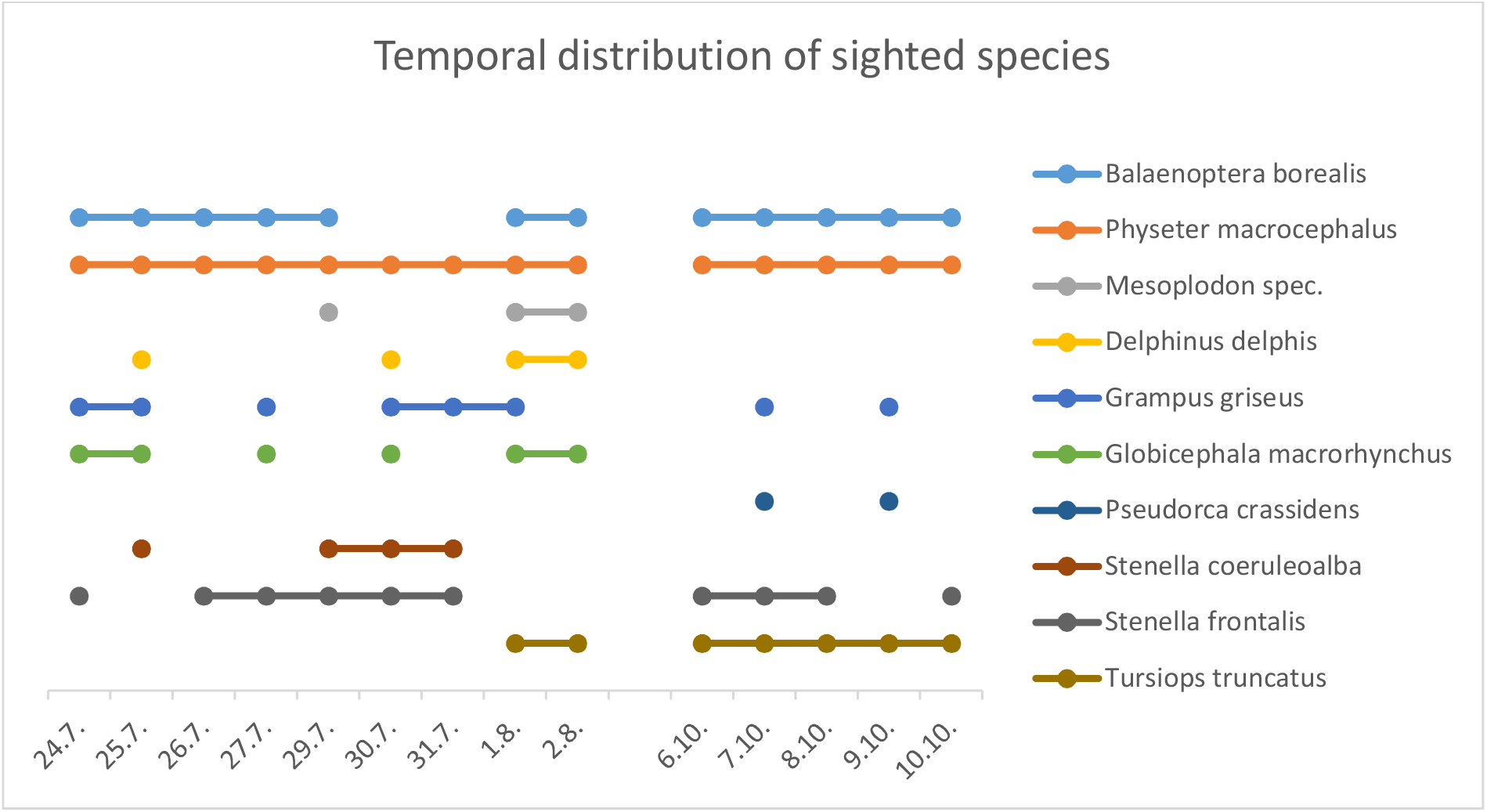
Temporal distribution of sighted species in July/August and October 2020.

The Sperm whale was the most observed cetacean species throughout the excursions in 2020 with 58 individuals registered. In July and August 42 individuals were recorded in nine days and in October 16 individuals in five days (Fig. 1).

Table 3 shows the sightings per day. Vieira and Brito (2009) wanted to know if a whaler or a whale-watcher was able to see more Sperm whales. The whaling and the whale-watching periods were examined. They obtained 593 sightings of *Physeter macrocephalus* during a total of 191 days of whaling from the years 1948, 1968, 1969, 1972, and 1973. Sightings are considered as both, effective captures as well as strikes that did not result in capture. Than data for 1133 days of whale-watching activities were examined, available information from Espaço Talassa between the years 1997 and 2008, with 1767 Sperm whales sightings. Their finding was that whaler observed more Sperm whale individuals than the whale watchers (Vieira and Brito, 2009). Furthermore table 3 shows the sightings per day for all excursions from the University of Hildesheim from the years 2016 to 2020.

**Tab.3:** Whaling and whale-watching sightings per day.

| Activity | Period | Sighted Sperm whales | Days | Sightings /day | References |
| --- | --- | --- | --- | --- | --- |
| Whale hunting | 1948-1973 | 593 | 191 | 3.1 | Vieira & Brito (2009) |
| Whale-watching | 1997-2008 | 1767 | 1133 | 1.6 | Vieira & Brito (2009) |
| Whale-watching | Excursion 2016 | 4 | 6 | 0.7 | P. Zahn |
| Whale-watching | Excursion 2017 | 29 | 11 | 2.6 | P. Zahn |
| Whale-watching | Excursion 2018 | 62 | 19 | 3.3 | P. Zahn |
| Whale-watching | Excursion 2019 | 106 | 17 | 6.2 | P. Zahn |
| Whale-watching | Excursion 2020 | 58 | 14 | 4.1 | P. Zahn |

## Discussion

Cetaceans are under increasing pressure from anthropogenic activities (Silva et al., 2014). In response to direct effects of climate change and anthropogenic-driven reorganization to their ecosystem cetaceans are expected to experience important changes in their distribution. Global climate change expose them to an altered marine environment (Bron et al., 2019). The Azores offer a high diversity of cetaceans. But information of spatial and temporal distribution of this marine megafauna group is still limited (Tobeña et al., 2016). The knowledge of the full geographic range of most cetacean species within the North Atlantic is restricted as well as the understanding of their seasonal movements within this range (Silva et al., 2014). This requires quantitative information on their spatial and temporal presence (Silva et al., 2014; Pérez-Jorge et al., 2020). This study attempted to gain useful opportunistic data for cetacean research.

In this study, members of 4 different families were recorded. This is in agreement with Bron et al. (2019), Silva et al. (2003) and Tobeña et al. (2016) who recorded members of this 4 families. In a much longer survey (10 years), Silva et al. (2014) observed members of 6 families.

During this 14 days short study a total of 10 whale and dolphins species were recorded. The mean number of sighted whale species per boat survey was 3.9. The only land-based observation showed 3 different species. This is remarkable, because of the short space of time and it demonstrates that the Azores are one of the best regions worldwide for whale-watching and cetacean research. It is consistent with the results from Silva et al. (2003). They recorded 11 different species during the surveys conducted in the summer and autumn seasons of 1999 and 2000. In a long term investigation with boat-surveys and land-based observations from 1999 to 2009 they sighted 24 different species (Silva et al., 2014). Tobeña et al. (2016) recorded 16 different species by using data collected by the Azores Fisheries Observer Programme between 2004 and 2009. Bron et al. (2019) used data collected by Futurismo Azores Adventures (São Miguel) with a total of 22 different cetacean species. With all species considered together the area shows a relative species richness.

### Balaenoptera borealis

Baleen whales are highly migratory marine species. This marine animals are increasingly vulnerable to population declines and extinction due to cumulative anthropogenic impacts on their environment. Understanding the distribution patterns are critical for effective conservation management (Pérez-Jorge et al., 2020) The annual migrations of baleen whales between low-latitude wintering grounds and high-latitude summer feeding areas leads to their presence at middle-latitude around the Azores at spring and summer (Pérez-Jorge et al., 2020; Silva et al., 2014; Still et al., 2019). Six species of the family Balaenopteridae may be seen in Azorean waters. Silva et al. (2014) reported that only Blue, Fin and Sei whale are sighted frequently every year, especially in spring and summer.

Visser et al., (2011a) showed that the time window, in which large baleen whales are at the Azores, depends more on the availability of prey than on the current time in the year. The timing of their presence is strongly related to the onset of the North Atlantic phytoplankton spring bloom (Visser et al., 2011a). And zooplankton development often tracks the phytoplankton spring bloom. Food availability therefore plays a key role in the spatial and temporal distribution. This is consistent with the results of Pérez-Jorge et al. (2020).

Tobeña et al. (2016) also explain a very variable spatio-temporal pattern of this group. This may be the reason that in this study only one representative of the suborder Mysticeti was sighted. The Sei whale is a member of the family of rorquals (Balaenopteridae). The number of sightings (20) for this species was unusually high and 35 individuals were seen. In July and August *Balaenoptera borealis* was seen 26 times (14 boat surveys) and 9 records (8 boat surveys) in October. The result is not in agreement with Silva et al. (2014) and Pérez-Jorge et al. (2020). This species is normally observed from March to July in the Azores and reports of their presence after August are rare (Pérez-Jorge et al., 2020; Silva et al., 2014). According to Visser et al. (2011a) low primary production in summer and autumn is the reason why large baleen whales avoid the Azores during their migration to the southern wintering areas in autumn. During their study no baleen whale species was observed between October and December. The frequent occurrence of *Balaenoptera borealis* in October 2020 recorded in this study therefore leads to the assumption that food availability was unusually good this autumn (Visser et al. 2011a; Pérez-Jorge et al., 2020).

### Physeter macrocephalus

The Sperm whale is the first of nine sighted species of Toothed whales in this study to discuss. *Physeter macrocephalus* is the only representative of the family Physeteridae. The region around the Azores seems to be an important habitat for species that typically inhabit deep waters. That is also true for the Sperm whale. They are considered to be essentially teutophagus. Individuals of this species are frequently observed in the Azores all year round. They are among the most frequently sighted species around all island of the archipelago and the main target of the whale-watching activities (Bron et al., 2019; Silva et al., 2003 und 2014; Tobeña et al., 2016). In this study, the Sperm whale has the highest number of sightings (30) of all observed whale species.

The females are resident while the adult males migrate throughout the year (Espaço Talassa 2020, Still et al. 2019). Silva et al. (2014) reported 76 % of the sightings comprised groups of adult females, subadults and calves, and 8 % were adult males which this study agrees with. This survey included 40 adult females, subadults and calves in July/August recorded and 15 in October, with 2 and 1 adult males respectively. From this follows that 5 % of the sightings are adult males (4.8 % in July/August and 6.3 % in October). Female groups often concentrate in relative small areas, where they form feeding aggregations (Silva et al., 2014), an observation which is confirmed by this study.

Land based whaling in the Azores archipelago ceased around 1987 after long decades. It was always conducted with open boats and employing hand-held harpoons and lances. The slow swimming and abundant Sperm whale was the only one to catch (Vieira and Brito, 2009). A few years after the end of whaling whale-watching begun, as an attempt to keep using a valuable natural resource and the need to employ former whalers and lookouts. Regarding the Azores the question arose, if a tourist is seeing fewer Sperm whales presently than a whaler could kill in the past (Vieira and Brito, 2009). Twenty years after the end of whaling they compared the number of sightings during whaling and whale-watching activities.

They showed, that fewer Sperm whale were sighted in the whale-watching activities 20 years after the end of whaling compared to the whaling years. In fact sightings in the past were significantly higher. Whalers sighted 3.1 Sperm whales per day (table 3) in comparison to 1.6 sightings per day (mean number from 1997 to 2008) for whale watchers. The total number of whaling sightings may be underestimated, as there may have been more individuals been sighted than those captured. Nevertheless, captures in the Azores greatly decreased the number of Sperm whales throughout the decades. Twenty years after the end of whaling a reduced number of recent sightings were found.

In this study, 58 individuals of *Physeter macrocephalus* were sighted in 22 days, which are 4.1 sightings per day. Table 3 also shows the results of former excursions from 2016 to 2019. Since 2018 the results exceeds the 3.1 sightings per day of whalers (Vieira & Brito, 2009), from 3.3 to 6.2 sightings per day during the excursions. Unlike Vieira and Brito (2009) this study valued every day of the excursions, including days without sightings of Sperm whales. It is therefore assumed, that a recovery of the population size took place 30 year after the end of whaling. It is also possible that 20 to 30 years after the end of whaling some more individuals of *Physeter macrocephalus* stopped avoiding the feeding grounds environment of the Azores, since they slowly learned that there is no more danger for them.

### Mesoplodon bidens

Silva et al. (2014) reported an almost year-round presence of Mesoplodon species with a peak in summer month and a fairly common presence. The excursions 2020 results are contrary. With three sightings and 14 individuals the record is low in this study. Only one sighting in July showed Sowerby’s beaked whale. The other two sightings in August were of the genus Mesoplodon but the identification of the species was not possible. The Azores, as the southern part of the range of *Mesoplodon bidens*, may represent a critical habitat for the latter (Silva et al., 2014). Tobeña et al. (2016) rather found an improvement of habitat suitability for beaked whales.

Silva et al. (2014) described the study area with a wide range of habitat types, including areas of abyssal plain, narrow island shelves, submarine canyons, steep island slopes and shallow seamounts. The particular bottom topography of the Azores conforms to the deep-water preference of Beaked whales (Silva et al. (2003). The deep diving species in the region, Sperm whale, Short-finned pilot whale, Risso’s dolpin, and Beaked whales, are considered to be teutophagus (Tobeña et al., 2016). In the Azores Sowerby’s beaked whale has a different diet, composed mostly of meso- and bathy-pelagic fish. This resource is abundant here also.

### Delphinus delphis

Some members of the family of Oceanic dolphins (Delphinidae) were the most dominant species for the entire survey. Of the seven sighted species *Delphinus delphis* is the first to discuss. Short-beaked common dolphin, Striped dolphin, Risso’s dolphin, and Common bottlenose dolphin are present in the Azores throughout the year (Silva et al., 2014; Tobeña et al., 2016). They and Bron et al. (2019) reported, that *Delphinus delphis*, and *Tursiops truncatus* were most frequently seen and *Grampus griseus* was frequently observed.

In this study, Short-beaked common dolphin was sighted 6 times. This is a low number in comparison to Risso’s dolphin (12), Common bottlenose dolphin (12), and Atlantic spotted dolphin (17). The result for *Delphinus delphis* is on the contrary with Silva et al. (2014) and Tobeña et al. (2016) who found Short-beaked common dolphin as the most often observed species. A probable explication are the survey period. In July, August and October the study was late in the year.

There is a succession pattern for *Delphinus delphis* and *Stenella frontalis* for their seasonality (Tobeña et al., 2016). With the occurrence of *Stenella frontalis* in the Azores coincides the displacement of *Delphinus delphis* when sightings of the first outnumbers those of *Delphinus delphis* (Silva et al., 2014). During that time from June to November the encounter rate of *Delphinus delphis* decreases (Silva et al., 2014). This is consistent with the result of this study. Tobeña et al. (2016) reported that the distribution of the Short-beaked common dolphin becomes restricted to some seamount complexes as the season progressed. That indicates, that the seamounts may play an important role in maintaining conditions for the occurrence of the species in the region throughout the year.

### Globicephala macrorhynchus

In addition to the smaller dolphins, also larger members of this family occur in the waters of the Azores. These include Short-finned pilot whale and False killer whale. According to the study by Silva et al. (2014), this two species can be spotted year round. The prediction for the two species varies substantially (Tobeña et al., 2016). Both species mainly visit the Azores in the summer months until autumn, which indicates a preference for higher water temperatures (Still et al., 2019). Sightings in winter (February) are also possible, as Silva et al. (2014) have proved.

Short-finned pilot whales and Risso’s dolphins are more similar to each other based on their feeding ecology (Tobeña et al., 2016). *Globicephala macrorhynchus* is not resident in the waters of the Azores (Still et al., 2019). They are regular visitors to the Azores and often seen from April to November (Silva et al., 2014). In this study the species was only observed in July / August in 2020 but was unusually frequent (8 sightings in 6 days). Silva et al. (2014) report still common sightings in October, but they become rarer the colder it gets. The results of this study does not confirm this, but it acknowledges the great variation of the encounter rate between month and years. The Azores lie on the northern limit of the Short-finned pilot whale distribution, the species is resident in warm subtropical waters and therefore avoids the Azores in the coldest months from December to March (Still et al., 2019).

### Grampus griseus

Risso’s dolphin and Short-finned pilot whale are both teutophagous and deep divers. *Grampus griseus* together with *Tursiops truncatus* is one of the most frequently sighted species in this area and can be sighted all year round (Silva et al., 2014; Still et al., 2019; Tobeña et al., 2016). The encounter rate of Risso’s dolphin varies greatly between months but they are among the five cetacean species with higher sightings rates (Silva et al., 2014). This study shows 12 sightings (9 in July/August and 3 in October) for *Grampus griseus*, second most after *Stenella frontalis*. This confirms both findings, the higher encounter rate for Risso’s dolphin and that they are frequently observed. Hartman et al. (2009) concluded a resident population of Risso’s dolphin in the Azorean waters. They favour habitats over steep slopes with depth between 400 and 1.200 m, preferably areas where the slope is close to the coastline (Pereira, 2008). Silva et al. (2003) reported an even distribution throughout the whole Archipelago.

### Pseudorca crassidens

Sightings of False killer whale are a rarity compared to the other sighted Delphinids. *Pseudorca crassidens* are not resident in the waters of the Azores (Still et al., 2019). A long-term study of Silva et al. (2014) showed that False killer whale regularly passes the Azores, but recorded a less obvious pattern of occurrence for this species. Due to their sporadic or rare incursions to the area, or because of their cryptic habits they are difficult to find at sea.

In 2020, the statistic from Espaço Talassa shows this species only three times earlier this year, two in July and one in September (Espaço Talassa 2020). More than a handful of sightings per year are uncommon. In this study the sightings in October were extraordinary, considering that the majority of sightings usually occur in July and August. The four sightings on October 7^th^ and 9^th^ suggest that it was the same pod, because of the recognizable different markings of individuals.

### Stenella coeruleoalba

Silva et al. (2014) report a nearly continuous presence of *Stenella coeruleoalba* in the Azores. The encounter rate is higher between May and July. This confirms the result of this study. Striped dolphin was seen four times in July, but no sighting in October. Their distribution is strongly influenced by water temperature (Tobeña et al., 2016). This might explain the higher encounter rate in mid-summer.

### Stenella frontalis

Silva et al. (2014) and Tobeña et al. (2016) found that *Stenella frontalis* was within the most commonly encountered species. The seasonality of the Atlantic spotted dolphin is very noticeable, with first sightings in May. Relative abundance is reached in July and August and by October the species disappear. In this study, *Stenella frontalis* exhibits the highest number of schools (17) within the Oceanic dolphins, which makes it the most abundant and dominant species in this survey. Especially on October 7^th^ with an observed superpod of more than 3000 individuals approximately. This school was an aggregation to prepare their southward journey back to warmer waters, a behavior explained by the skippers (personal conversation). Silva et al. (2014) reported a similar aggregation observed in autumn 2008. That year they saw an exceptionally high number of individuals of *Stenella frontalis* in October as well. This result confirms also Silva et al. (2003) and Bron et al. (2019) with their findings that Atlantic spotted dolphin registered the highest sighting rate and was the most abundant species.

### Tursiops truncatus

Common bottlenose dolphin sightings are concentrated in the coastal area around the central group of the islands (Silva et al., 2003 und 2014; Tobeña et al., 2016; Bron et al., 2019). *Tursiops truncatus* is typically found over the island shelves and slopes and around shallow banks, in < 700 m deep areas. There is a considerable overlap in spatial distribution.

Common bottlenose dolphin are among the most frequently sighted species and occur year-round (Silva et al., 2014). But encounter rates varies greatly between months. This confirms the results with four observations in August and an especially high encounter rate in October (eight sightings). This represents a very different encounter rate between July/August and October. Tobeña et al. (2016) also described a great heterogeneity in distribution with Common bottlenose dolphin among different month which may reflect a contrasting influence of oceanographic processes on the distribution of cetacean species. Temporary immigration of large numbers of non-resident dolphins into the study area could explain the fluctuations in the number of sightings per month (Silva et al., 2014).

## Conclusions

The International Whaling Commission (IWC) describes in its whale-watching handbook the common research methods used to study whales and dolphins. They recommend a few of these methods to be easily combined with whale-watching. One of these are boat surveys to study distribution and habitat use (IWC, 2020). The knowledge obtained here could be used for policies aimed at effectively protecting cetaceans and their habits or in the development of management plans for specific areas, e.g. the definition of a load capacity for whale-watching activities (Silva et al., 2003; Tobeña et al., 2016). Whale-watching tours are often offered year-round and the data collected may be a potential tool for detecting long term changes (Bron et al., 2019; Silva et al., 2014).

This study illustrates the potential of using alternative sources of information to obtain meaningful data. Data used in this research were collected as a by-product of whale-watching, which was the primary purpose. This method is subject to various limitations. The normal tourism activity favours the observation of different species (Pereira, 2008). Therefore, the study area may not be equally covered. Coincidence may play an important role. Findings from this research may not be representative for the entire research area. The boat surveys during the excursions were not designed for the purpose to estimating cetacean abundance (Bron et al., 2019; Silva et al., 2014).

Nevertheless, the activity of whale-watching offer a source of valuable data. It represents a cost-effective method to collect opportunistic data on the cetaceans. Otherwise the information may be inaccessible, e.g. for rare species or incidental sightings. This study obtained opportunistic data that may be helpful to a better understanding of the temporal distribution of the cetacean species sighted.

## Acknowledgments

Sincere thanks are given to the staff of Espaço Talassa for their friendly support in all matters of the preparation and execution of the excursion. We are especially grateful to skippers and lookouts for their kindness, their endless passionate dedication to cetacean observation, and their enormous readiness for environmental education. We acknowledge the help of the students throughout this study, their commitment and enthusiasm for the protection of whales.

